# Thioredoxin reductase 3 inhibition sensitizes triple-negative breast cancer cells to EGFR inhibitors

**DOI:** 10.1101/2025.03.16.642742

**Authors:** Prahlad V Raninga, Göknur Giner, Sivanandhini Sankarasubramanian, George Ambalathingal, Murugan Kalimutho, Marco J Herold, Kum Kum Khanna

## Abstract

Triple-negative breast cancer (TNBC) is an aggressive subtype with poor prognosis and limited treatments. Although 40-70% of TNBC cases overexpress EGFR, anti-EGFR therapies show minimal clinical benefit. This may result from inherent resistance, inactive EGFR, or its absence on the plasma membrane. Here, we used genome wide CRISPR knock out library screening to identify factors that mediate resistance to EGFR inhibitor. We discovered that depletion of a redox protein, thioredoxin reductase 3 (TXNRD3), in MDA-MB-231 cells reduces cell survival after Erlotinib treatment suggesting that loss of TXNRD3 may sensitize TNBC cells to EGFR inhibitors. siRNA-induced knockdown or pharmacological inhibition of TXNRD3 using an FDA-approved drug Auranofin significantly sensitized EGFR-high TNBC cells to EGFR inhibitors. Mechanistically, TXNRD3 knockdown or inhibition using Auranofin increased oxidative stress-mediated accumulation of phosphorylated EGFR at Y1068 and increased surface accumulation. Interestingly, combination of Auranofin with EGFR inhibition markedly induced antibody-dependent cell mediated cytotoxicity (ADCC) and exerted a significant anti-cancer activity *in vivo* model. Overall, our findings indicate that targeting TXNRD3 with Auranofin can activate EGFR and enhance its surface localization, thereby sensitizing TNBC cells to anti-EGFR therapies. This approach offers a promising strategy for treating TNBC patients who are resistant to current EGFR-targeted treatments.

## BACKGROUND

Triple-negative breast cancers (TNBCs) are most aggressive subtype with a high histology grade, and account for approximately 15-20% of all diagnosed invasive breast cancer cases world-wide and are more prevalent in younger women (<40 years of age). TNBCs are highly heterogeneous and can be subdivided into four types including basal-like, luminal, mesenchymal, and immune-modulatory/activated (1). Due to the lack of surface hormone receptors, TNBCs cannot be treated by anti-hormonal therapies unlike luminal and HER2+ breast cancers. Hence, chemotherapies and radio therapy remain mainstay treatment for TNBC patients. Although a majority of TNBC patients respond to initial chemotherapies, these tumors are prone to recur, metastasize (disseminate to the lungs and brain), and acquire chemoresistance (2). Unfortunately, the severe adverse effects and drug resistance associated with standard cytotoxic chemotherapies (e.g. anthracyclines, cyclophosphamide, and taxanes) compromises their clinical benefits in TNBC treatment (3,4). Thus, there is an urgent need to explore clinically viable targeted therapies for primary and metastatic TNBCs.

Several studies have reported EGFR overexpression in 40-70% of TNBC patients by immunohistochemistry (5,6). EGFR overexpression correlates with poor clinical outcome and decreased disease-free survival in TNBC patients (5). Elevation of EGFR messenger RNA (mRNA), as well as activation of the EGFR pathway in gene expression microarray studies, is a characteristic of basal-like breast cancer (7,8), and is associated with poor prognostic signatures in almost all basal-like tumors (9). Thus, EGFR represents a clinically relevant target in basal-like TNBC patients. Based on these findings, a small molecule EGFR inhibitor, Erlotinib, has been tested in the clinical trials in TNBC patients either as monotherapy or combination therapies. Results from Erlotinib monotherapy in unselected previously treated women with locally advanced or metastatic breast cancer metastatic and recurrent TNBC showed minimal activity (10). Similarly, phase II trial of an EGFR monoclonal antibody cetuximab in combination with carboplatin in stage IV TNBC patients showed no significant clinical response (11). Hence, despite EGFR overexpression representing an attractive therapeutic target, EGFR inhibitors failed to exert a significant clinical activity potentially due to the existence of inherent resistance to the drug. Understanding such mechanisms and their simultaneous inhibition may sensitize TNBC cells to EGFR inhibitors.

In this study, we have identified potential drivers of EGFR inhibitor response in basal-like TNBCs. We performed a genome wide CRISPR/Cas9 knockout screening in TNBC cells with Erlotinib treatment to systematically evaluate the driving mechanisms of Erlotinib resistance. We identified thioredoxin reductase 3 (*TXNRD3)*, a member of the thioredoxin protein family involved in maintaining intracellular redox homeostasis, as a critical driver of Erlotinib resistance in TNBC cells. We found that genetic depletion of *TXNRD3* sensitized TNBC cells to EGFR inhibitors, Erlotinib and Osimertinib. Additionally, a recent study showed that *NRF2*-driven antioxidant genes including TXNRDs, are upregulated in Osimertinib-treated drug persister lung cancer cells (12) suggesting the potential involvement of thioredoxin genes in EGFR inhibitor resistance. Interestingly, co-inhibition of TXNRD3 using an FDA-approved thioredoxin inhibitor Auranofin sensitized basal-like TNBC cells to EGFR inhibitors *in vitro* and *in vivo*.

## MATERIALS & METHODS

### Cell lines, reagents, and patient tissue samples

All breast cancer cell lines were obtained from the American Type Culture Collection (ATCC). The SUM159PT cell line was kindly provided by Prof Sunil Lakhani, UQCCR, Australia. The 4T1.2 cell line was kindly provided by Dr Robin Anderson, Olivia Newton-John Cancer Research Institute, Australia. CRISPR/Cas9 EGFR knockout (KO) and CRISPR/Cas9 Vector control MDA-MB-231 cells were generated by Prof Chia-Hwa Lee (Taipei Medical University, Taiwan) (13). All breast cancer cell lines were cultured in DMEM media supplemented with 10% fetal bovine serum (FBS) and 100 nM Sodium Selenite. All cell lines were tested for Mycoplasma infection and authenticated using short tandem repeat (STR) profiling by scientific services at QIMR Berghofer Medical Research Institute. Auranofin was purchased from Cayman Chemicals (Cat #: 15316). Erlotinib (Cat #: S7786) and Osimertinib (Cat #: S7297) were purchased from Selleck Chemicals. TRi-1 was purchased from MedChem Express (Cat #: HY-125006. The list of antibodies used in this study is provided in supplementary Table S1. TNBC patient tissue microarray was provided by the Mater Pathology and Mater Cancer Hospital, Brisbane.

### Cell viability assays

EGFR-high breast cancer cells (SUM159PT, MDA-MB-231, HCC1806, and HCC1143) were seeded at the density of 3000 cells/well onto a white-walled clear bottom 96-wells plate overnight. Cells were then treated with Auranofin and EGFR inhibitors, Erlotinib or Osimertinib, alone or in combination for 3 days, and cell viability was analyzed using the Real-Time-Glo™ MT Cell Viability Assay (Promega) as per the manufacturer’s guidelines. Synergy scores between different drug combination was calculated using Synergy Finder (14). To analyze the effect of Auranofin and Erlotinib combination therapy on EGFR-low breast cancer cells, MCF7, BT-474, and SKBR3 cells were seeded at the density of 3000 cells/well onto a clear 96-wells plate. Cells were treated with Auranofin and Erlotinib, alone or in combination, for 3 days, and cell growth was analyzed by MTS assays (Promega).

### Genome-wide CRISPR/Cas9 knockout library screen

In this study, we used the Human GeCKOv2A & 2B CRISPR knockout pooled library to identify genes responsible for Erlotinib resistance in TNBC cells. The library was a gift from Feng Zhang (Addgene # 1000000049) (15). The workflow of this forward CRISPR-based genetic screen is illustrated in Fig 1A. Firstly, we established a stable Cas9-expressing MDA-MB-231 cell line by lentiviral transduction of Cas9 coding sequence. We then transduced Cas9-expressing MDA-MB-231 cells with GeCKO v2A and B library which contains 123,411 unique sgRNA sequences targeting 19,052 human genes and 1864 miRNAs (6 sgRNAs per gene, 4 sgRNAs per miRNA, and 1000 non-targeting control sgRNAs) at an MOI 0.3 to ensure each cell represents one sgRNA. The transduced cells were then selected with 1 µg/mL of puromycin for 7 days to generate a mutant cell line. The mutant cells were then treated with vehicle (DMSO) and Erlotinib (10 µM) for 7 days, respectively. After 7 days of treatment, the residual cells were collected, and genomic DNA was isolated from each treatment group. The sgRNA sequences were amplified and were subjected to parallel amplicon sequencing at Ramaciotti Centre for Genomics to identify sgRNAs lost in Erlotinib-treated cells.

**Figure 1:**
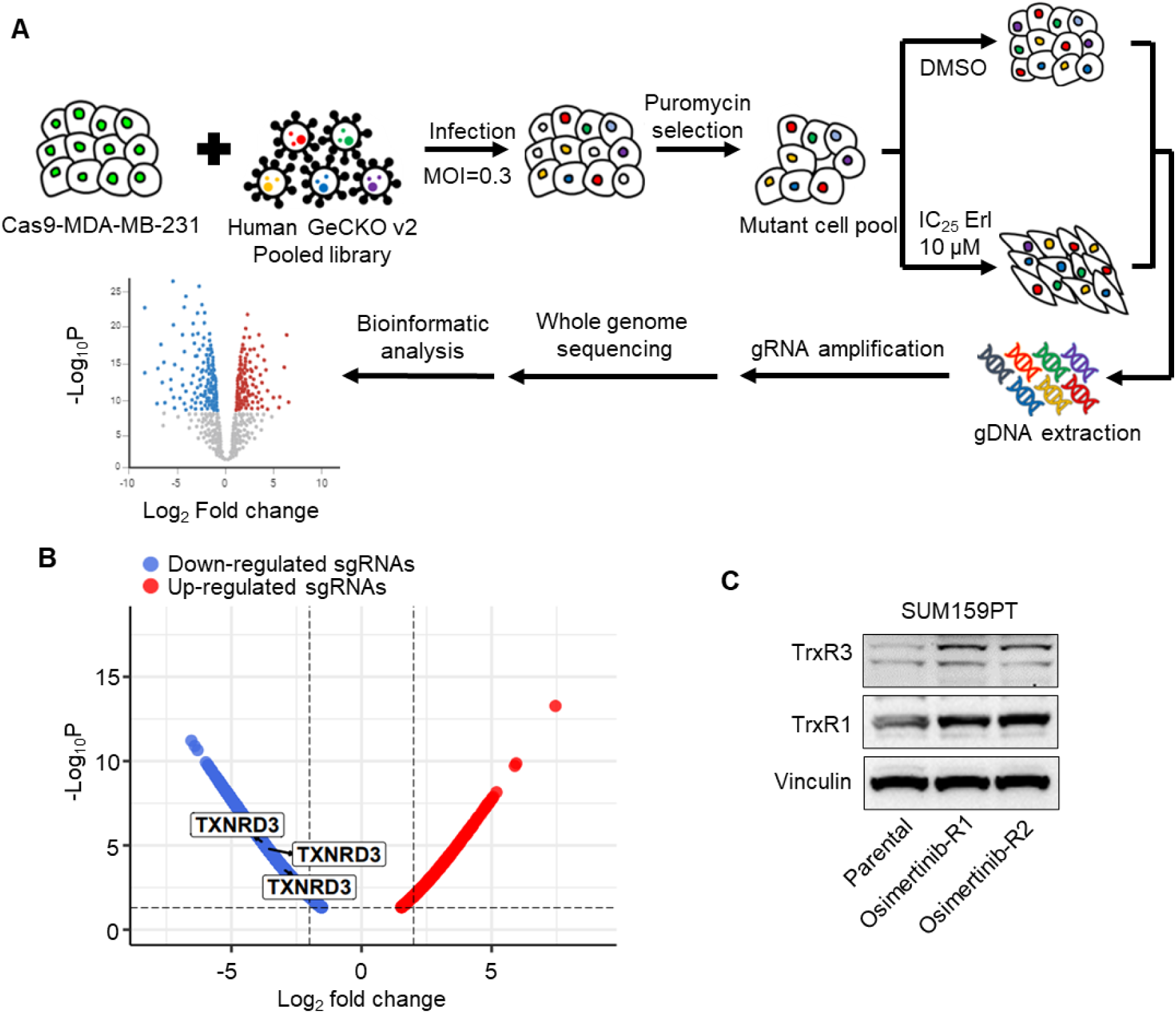
CRISPR library screening identified *TXNRD3* as a critical gene for Erlotinib resistance in TNBC cells. **(A)** Schematic diagram illustrating the workflow of genome wide CRISPR/Cas9 knockout library screening (CRISPR: Clustered Regularly Interspaced Short Palindromic Repeats) of Cas9-MDA-MB-231 cells treated with either DMSO or Erlotinib (IC_25_, 10 µM). Figure was adapted (38) and modified using BioRender. **(B)** Volcano plot illustrating the differential guides significantly depleted (in blue) or enriched (in red) in Erlotinib-treated MDA-MB-231 cells compared to DMSO-treated control MDA-MB-231 cells. The differential guides were plotted statistical significance (-log_10_ p-value) on the y-axis against fold change (log_2_) on the x-axis. Each point represents a guide, with those above the significance threshold with fold change greater than 2. TXNRD3 guides are highlighted. Source data are provided as a supplementary source file. **(C)** Protein levels of thioredoxin reductase 1 (TrxR1) and thioredoxin reductase 3 (TrxR3) were analyzed by Western blot analysis in parental and Osimertinib-resistant or persister SUM159PT cells. Representative images of three independent Western blot analysis are shown. Vinculin was used as a loading control.

### CRISPR data analysis

#### Exploratory Analysis of sgRNA Counts

For the exploration analysis of sgRNA counts, we employed several visualization techniques using R packages. Pie charts were generated using the ‘scales‘ package (version 1.3.0) to provide an overview of the distribution of sgRNA counts. Volcano plots, depicting the fold change and statistical significance of differential expression, were created using the ‘Enhanced Volcanò package (version 1.20.0). Additionally, waterfall plots were generated using the ‘waterfalls‘ package (version 1.0.0) to visualize changes in sgRNA abundance across different conditions.

#### Differential Abundance Analysis

Differential abundance analysis was conducted using the ‘edgeR‘ Bioconductor package (version 4.0.12) (16,17). Initially, lowly expressed genes were filtered out based on criteria of having a count lower than the median count per library for at least two samples. Subsequently, the counts were normalized using the upper-quartile method (18), implemented at the 0.99 quantile within the ‘normLibSizes‘ function.

Differential abundance of sgRNA counts between treatments was assessed using gene wise likelihood ratio tests within the ‘edgeR‘ framework. To control for multiple testing, resulting p-values were adjusted using the false discovery rate (FDR) method of Benjamini and Hochberg (19).

### TXNRD3 ELISA assays

SUM159PT and MDA-MB-231 cells (5 x 10^5^ cell) were seeded in 60 mm dish overnight and treated with Auranofin (0-2.5 µM) for 24 hours. Then, cells were collected, and protein was extracted. TXNRD3 levels were analyzed in the protein lysates using Human Thioredoxin Reductase 3 (TXNRD3) ELISA Kit (abx384053, Abbexa, UK) as per the manufacturer’s instructions.

### siRNA transfections

SUM159PT and MDA-MB-231 cells were transfected with 20 nM of non-specific scramble small interfering RNAs (siRNAs), human TrxR1-specific, human TrxR3-specific, OR human EGFR-specific siRNAs using Lipofectamine RNAi MAX (Invitrogen Cat #: 13778030) per manufacturer’s instructions. The sequences of siRNAs used in manuscript are provided in supplementary table S2.

### Intracellular ROS assays

Breast cancer cell line, SUM159PT and MDA-MB-231, were seeded overnight (5000 cells/well) in black-walled 96-well plates and were treated with or without Auranofin for 24 h. The media containing drug was then removed, cells were washed with 1X PBS and incubated with 5 µM H2DCFDA for 30 mins at 37^0^C, and fluorescence was measured using a H4 Synergy H4 Multi Mode Plate Reader (Syn Biotek).

### Thioredoxin reductase 1 activity assays

Breast cancer cells were treated with Auranofin for 24 h, then lysed, and cellular TrxR1 redox activity was measured by the DTNB reduction assay as described previously (20).

### Immunofluorescence

SUM159PT (3 x 10^5^) cells were seeded on glass coverslips and allowed to adhere to the surface overnight. Cells were then treated with or without Auranofin (2.5 µM) for 24 hours. The coverslips with cells were fixed using 4% paraformaldehyde (PFA) (Sigma Aldrich, USA) for 15 minutes at room temperature (RT). Subsequently, coverslips were blocked in a solution of 2% Bovine Serum Albumin (BSA) (Sigma Aldrich, USA) in PBS for 1 hour at RT. EGFR antibody (dilution 1:200) were added to the blocking buffer, and coverslips were incubated with this antibody mixture for 1 hour at 37°C. Following this, the coverslips were washed three times with PBS and then incubated with Alexa-Fluor-conjugated secondary antibody (AF488) and DAPI (4′,6-diamidino-2-phenylindole) for nuclei staining in a 37°C incubator for 30-45 minutes. After incubation, coverslips were washed three times with PBS and mounted on a glass slide using mounting medium Prolong® Gold anti-fade from Life Technologies™. Images were captured using a Delta-Vision personal DV deconvolution microscope (Applied Precision, GE Healthcare, Issaquah, WA). The analysis of the images was performed using Fiji ImageJ software (National Institutes of Health, USA).

### Reverse Transcription-qPCR

SUM159PT and MDA-MB-231 cells were transfected with either scramble siRNA (siControl), TrxR1-specific siRNA, or TrxR3-specific siRNA for 48 hours. TrxR1 and TrxR3 gene expression analysis using reverse-transcriptase quantitative PCR (RT-qPCR) was performed using the Viia7 real-time PCR system (Applied Biosystems, US).

### Antibody-dependent cellular cytotoxicity (ADCC) assays

Real-time killing assay was performed using xCelligence MP platform (ACEA Biosciences). 96 well E-plates were seeded with 20000 SUMPT cells/well in 150 µl RPMI+10% FCS medium. The cells were allowed to attach and proliferate for 20 hours before being treated with Auranofin (1 µM), cetuximab (50 µg/ml), PBMCs (E:T 10:1), a combination of Auranofin (1 µM) + PBMCs (E:T 10:1), cetuximab (50 µg/ml) + PBMCs (E:T 10:1) and Auranofin (1 µM) + cetuximab (50 µg/ml) + PBMCs (E:T 10:1). Data normalization and analysis were performed using RTCA software (ACEA Biosciences). The PBMC cells were isolated from a healthy volunteer.

### Western blot

Breast cancer cells were either transfected with specific siRNAs as described above, or treated with Auranofin (0-2.5 µM, 24 hours). Proteins were extracted using a 7M Urea buffer. Immunoblotting was performed as described previously (21) with the antibodies listed in Table S1. The Super Signal chemiluminescent ECL-plus (Amersham) was applied for antibody detection.

#### *In vivo* animal models

All experiments were conducted in accordance with the guidelines of the QIMR Berghofer Medical Research Institute Animal Ethics Committee. For human MDA-MB-231 xenograft, 3 X 10^6^ MDA-MB-231 cells were prepared in 50% growth factor-reduced Matrigel (BD, Biosciences, Bedford, USA)/PBS and injected into the right 4^th^ inguinal mammary fat pad of 6-weeks old female Balb/c Nude mice. For murine 4T1.2 syngeneic models, 1 X 10^5^ cells prepared in 1X PBS were injected into the right 4^th^ inguinal mammary fat pad of 6-weeks old female immunocompetent Balb/c mice. For all *in vivo* mouse models, once tumor size reached ∼30-50 mm^3^, mice were randomized blindly into four treatment groups: (i) vehicle, (ii) Erlotinib (50 mg/kg, Oral, Monda-Friday), (iii) Auranofin (half-MTD, 5 mg/kg, ip, Monday-Friday), and (iv) combination for 3 weeks. Tumor growth was measured thrice weekly using a digital caliper. To calculate the tumor volume the following formula was used: tumor volume = [Lx W^2^]/2, where W = width of the tumor and L = length of the tumor.

### Statistical analysis

All values are presented as mean ± SEM. Data were analyzed using GraphPad Prism 9 (GraphPad Software, CA, USA). Statistical significance was determined by t-Test when comparing two groups only or by ANOVA for multiple comparisons. Specific post-tests were applied with ANOVA as indicated in each figure legend.

## RESULTS

### CRISPR library screening identified *TXNRD3* as a critical gene for Erlotinib resistance in TNBC cells

We performed genome-wide CRISPR/Cas9 knockout library screen to identify critical genes involved in Erlotinib resistance in human TNBCs. We transduced Cas9 expressing MDA-MB-231 cells that express high EGFR levels with human GeCKO v2 CRISPR lentiviral library (contains 123,411 unique sgRNAs, targeting 19,052 protein-coding genes, 1864 microRNAs, and 1000 control sgRNAs) at an MOI of 0.3 (Fig. 1A). After puromycin selection, we achieved about 400× coverage of the library and around 90% of the sgRNA sequences were retained in all samples (Fig. S1A), which ensure the sufficient read death and library coverage for the CRISPR library screening. The stably transfected cell pool was treated with vehicle or Erlotinib (IC25 - 10µM) for 7 days, genomic DNA was harvested and subjected to Next Generation Sequencing to identify sgRNA that are depleted upon Erlotinib treatment. We removed the non-targeting sgRNAs and guides with low expression levels to improve the quality of our library screening data and determined the library sizes in each treatment condition (Fig. S1B). We hypothesized that knockout of Erlotinib resistance driver genes will sensitize TNBC cells to Erlotinib-induced cell death or proliferation suppression. From this CRISPR/Cas9 knockout library screening, we identified a subset of sgRNAs targeting 1984 pan-cancer genes that were significantly depleted (P < 0.05) in the Erlotinib-treated cells when compared to vehicle control, indicating that these genes might be potential drivers for Erlotinib resistance (Fig. 1B h& Additional file 1). Amongst these 1984 depleted genes, 88% were targeted by >four sgRNAs, respectively (Fig. S1C). For validation studies, we focused on genes for which FDA-approved inhibitor is available. Among the list of genes, thioredoxin reductase 3 (*TXNRD3*), was identified as the most negatively selected gene upon Erlotinib treatment as cells expressing three different sgRNA specific to *TXNRD3* were lost after Erlotinib treatment (Fig. 1B).

*TXNRD3* is a member of the thioredoxin system. Three thioredoxin reductase isoforms are found in mammals, and are selenocysteine-containing flavoenzymes, which reduces thioredoxins and other cysteine-containing proteins, and plays a key role in redox homeostasis (22). *TXNRD3* is localized in the mitochondria and unlike other two isoforms, contains an additional N-terminal glutaredoxin domain allowing *TXNRD3* to participate in both thioredoxin and glutathione antioxidant systems.

In parallel with the CRISPR/Cas9 knockout library screening, we also examined TXNRD3 expression levels in EGFR inhibitor (EGFRi) Osimertinib-treated persister cells. We treated SUM159PT, a TNBC cell line with high EGFR expression, with an IC90 concentration of Osimertinib for 14 days and extracted protein from the residual Osimertinib-persister cells. TXNRD3 protein levels were significantly increased in Osimertinib-persister cells compared to parental (untreated) SUM159PT cells (Fig. 1C). TXNRD3 has been shown to be upregulated in TNBC cell lines as well as tumor tissues and is associated with adverse prognosis in TNBC patients (23). Moreover, higher TXNRD3 expression has been observed in Sorafenib-resistant leukemia and hepatocellular carcinomas (22). High TXNRD3 activity has been shown to maintain thioredoxin 2 in a reduced state, which in turn stabilizes several anti-apoptotic proteins, including Bcl-XL, Bcl-2, and MCL-1, to promote drug resistance (22), further suggesting the role of *TXNRD3* in conferring drug resistance. Together, our preliminary results and literature indicated that *TXNRD3* may play an important role in the development of EGFRi resistance in TNBCs.

### Genetic depletion of TXNRD3 sensitized TNBC cells to EGFR inhibitors

To further confirm the functional significance of TXNRD3 in conferring resistance to EGFR inhibitors and to determine whether TXNRD3 knockdown sensitizes TNBC cells to EGFR inhibition, we knocked down TXNRD3 expression using specific siRNAs in SUM159PT (Fig. 2A) and MDA-MB-231 cells (Fig. 2C). Additionally, we knocked down the expression of TXNRD1, a cytosolic isoform of thioredoxin reductase that has been shown to be upregulated in TNBC cells (24) to examine its potential role in EGFR inhibitor resistance (Fig. 2A & C). SUM159PT and MDA-MB-231 cells, transfected with either control siRNAs, TXNRD1-specific siRNAs, or TXNRD3-specific siRNAs, were treated with Erlotinib (0–10 µM) or Osimertinib (0–2.5 µM) for 72 hours. Cell viability was assessed using the MT cell viability assay. We found that TXNRD3 knockdown significantly sensitized both SUM159PT (Fig. 2B) and MDA-MB-231 (Fig. 2D) cells to both EGFR inhibitors, Erlotinib and Osimertinib. However, TXNRD1 knockdown did not sensitize SUM159PT or MDA-MB-231 cells to Erlotinib or Osimertinib (Fig. 2B, D). These findings suggest that TXNRD3 represents a promising therapeutic target for sensitizing TNBC cells to EGFR inhibitors.

**Figure 2:**
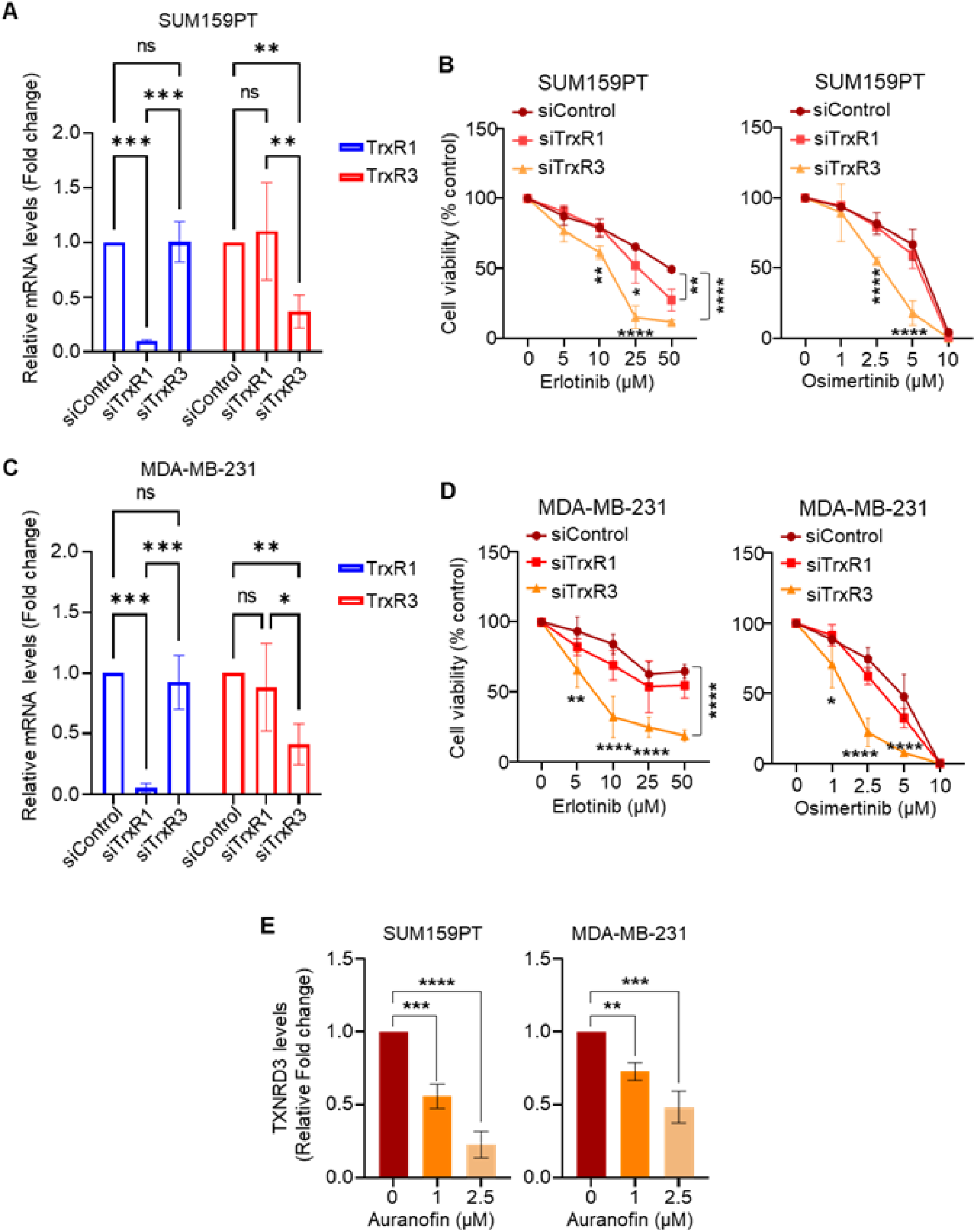
Genetic depletion of TXNRD3 sensitized TNBC cells to EGFR inhibitors. **(A)** SUM159PT cells were transfected either with scramble or control siRNA, TrxR1-specific siRNA, or TrxR3-specific siRNA for 48 hours. TrxR1 and TrxR3 mRNA levels were analyzed by RT-qPCR analysis. One-way ANOVA followed by Tukey’s post-test, n = 3 (mean ± SEM). **(B)** SUM159PT cells were transfected with either scramble or control siRNAs, TrxR1-specific siRNAs, or TrxR3-specific siRNAs for 24 hours and subsequently treated with Erlotinib (0-50 µM) or Osimertinib (0-10 µM) for 72 hours. Cell viability was analyzed by MT cell viability assays. One-way ANOVA followed by Tukey’s post-test, n = 3 (mean ± SEM). **(C)** MDA-MB-231 cells were transfected either with scramble or control siRNA, TrxR1-specific siRNA, or TrxR3-specific siRNA for 48 hours. TrxR1 and TrxR3 protein levels were analyzed by Western blot analysis. Representative images of three independent experiments are shown. GAPDH was used as a loading control. **(D)** MDA-MB-231 cells were transfected with either scramble or control siRNAs, TrxR1-specific siRNAs, or TrxR3-specific siRNAs for 24 hours and subsequently treated with Erlotinib (0-50 µM) or Osimertinib (0-10 µM) for 72 hours. Cell viability was analyzed by MT cell viability assays. One-way ANOVA followed by Tukey’s post-test, n = 3 (mean ± SEM). **(E)** SUM159PT and MDA-MB-231 cells were treated with Auranofin (0-2.5 µM) for 24 hours. Intracellular levels of TXNRD3 were analysed by ELISA assays. One-way ANOVA followed by Tukey’s post-test, n = 3 (mean ± SEM).

### Pharmacological inhibition of TXNRD3 using Auranofin sensitized TNBC cells to EGFR inhibitors *in vitro*

Since genetic depletion of TXNRD3 sensitized TNBC cells to EGFR inhibitors, we next examined whether pharmacological inhibition of TXNRD3 using an FDA-approved thioredoxin reductase inhibitor, Auranofin, could sensitize TNBC cells to EGFR inhibitors. We previously demonstrated that Auranofin inhibits TNBC tumor growth in vitro and in vivo by suppressing thioredoxin reductase redox activity (24). Notably, Auranofin has been shown to sensitize hepatocellular carcinoma cells to Sorafenib, a pan-receptor tyrosine kinase inhibitor (PDGFR, VEGFR, c-Kit, and Ret), through TXNRD3 inhibition (22). Based on this, we tested whether Auranofin inhibits TXNRD3 levels in TNBC cells and enhances their sensitivity to EGFR inhibitors. SUM159PT and MDA-MB-231 cells were treated with Auranofin (0–2.5 µM) for 24 hours, and TXNRD3 levels in cell lysates were analyzed using ELISA assays. We observed that Auranofin significantly reduced intracellular TXNRD3 levels in both SUM159PT and MDA-MB-231 cells in a concentration-dependent manner (Fig. 2E). Since

Auranofin significantly inhibited TXNRD3 in TNBC cells (Fig. 2E), we next examined whether Auranofin sensitizes TNBC cells to EGFR inhibitors. We treated four EGFR-high TNBC cell lines—SUM159PT, MDA-MB-231, HCC1806, and HCC1143—with a sub-lethal concentration of Auranofin (1 µM) in combination with EGFR inhibitors (Erlotinib and Osimertinib) and analyzed cell viability using MT assays. In line with previous studies, both Erlotinib and Osimertinib as single agents had a minimal effect on TNBC cell growth (Fig. 3A-D & Fig. S2A-D). However, co-treatment with Auranofin significantly sensitized SUM159PT, MDA-MB-231, HCC1806, and HCC1143 cells to both Erlotinib (Fig. 3A, B & Fig. S2A, B) and Osimertinib (Fig. 3C, D & Fig. S2C, D). Furthermore, we examined whether Auranofin exerts synergistic anti-cancer activity with EGFR inhibitors using the BLISS synergy score, where a score >10 indicates synergy, a score < -10 indicates antagonism, and a score between -10 and 10 indicates an additive effect. Our analysis revealed that Auranofin exhibited synergistic anti-cancer activity with both Erlotinib and Osimertinib in all four TNBC cell lines tested (Fig. 3A-D & Fig. S2A-D).

**Figure 3:**
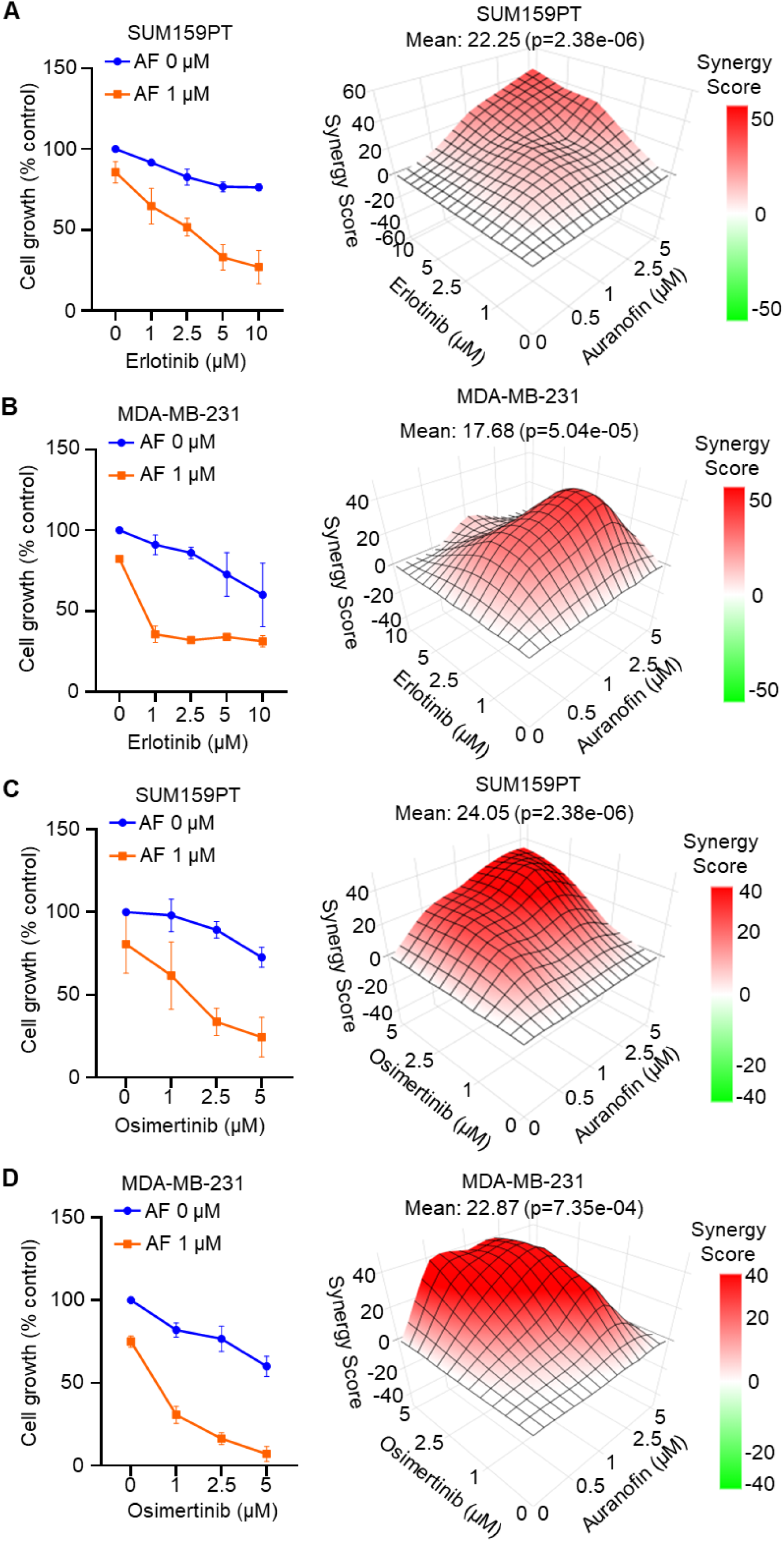
Pharmacological inhibition of TXNRD3 using Auranofin sensitized TNBC cells to EGFR inhibitors *in vitro*. **(A, B)** SUM159PT (A) and MDA-MB-231 (B) cells were treated with Auranofin (AF) (0-5 µM) and Erlotinib (0-50 µM), both alone and in combination, for 72 hours, and cell viability was analyzed by MT cell viability assays. For cell growth curves (left panel), results with only 1 µM AF are shown. Synergy score was calculated using Synergy Finder. One-way ANOVA followed by Tukey’s post-test, n = 3 (mean ± SEM). **(C, D)** SUM159PT (C) and MDA-MB-231 (D) cells were treated with Auranofin (0-5 µM) and Osimertinib (0-10 µM), both alone and in combination, for 72 hours, and cell viability was analyzed by MT cell viability assays. For cell growth curves (left panel), results with only 1 µM AF are shown. Synergy score was calculated using Synergy Finder. One-way ANOVA followed by Tukey’s post-test, n = 3 (mean ± SEM).

To further determine whether the synergistic anti-cancer activity between Auranofin and EGFR inhibitors is selective to EGFR-high breast cancer cells, we tested whether this combination affects EGFR knockout MDA-MB-231 cells. Our results showed that while Auranofin and EGFR inhibitors (Erlotinib and/or Osimertinib) significantly reduced the viability of vector control MDA-MB-231 cells, EGFR knockout MDA-MB-231 cells were partially resistant to Auranofin-Erlotinib (Fig. 4B) and Auranofin-Osimertinib (Fig. 4C) co-treatment. Additionally, siRNA-induced EGFR knockdown (Fig. 4D) rescued SUM159PT cells from Auranofin-Erlotinib combination therapy-induced cell death (Fig. 4E). We also tested whether Auranofin sensitizes EGFR-low breast cancer cells to Erlotinib. To this end, we treated EGFR-low breast cancer cell lines, including MCF7, BT-474, and SKBR3, with a sub-lethal concentration of Auranofin (1 µM) and Erlotinib (0–10 µM) for 72 hours and analyzed cell growth using MTS assays. Our results showed that Auranofin did not sensitize EGFR-low breast cancer cells to Erlotinib (Fig. S3), suggesting that the Auranofin-EGFR inhibitor synergistic anti-cancer effect is specific to EGFR-high TNBC cells.

**Figure 4:**
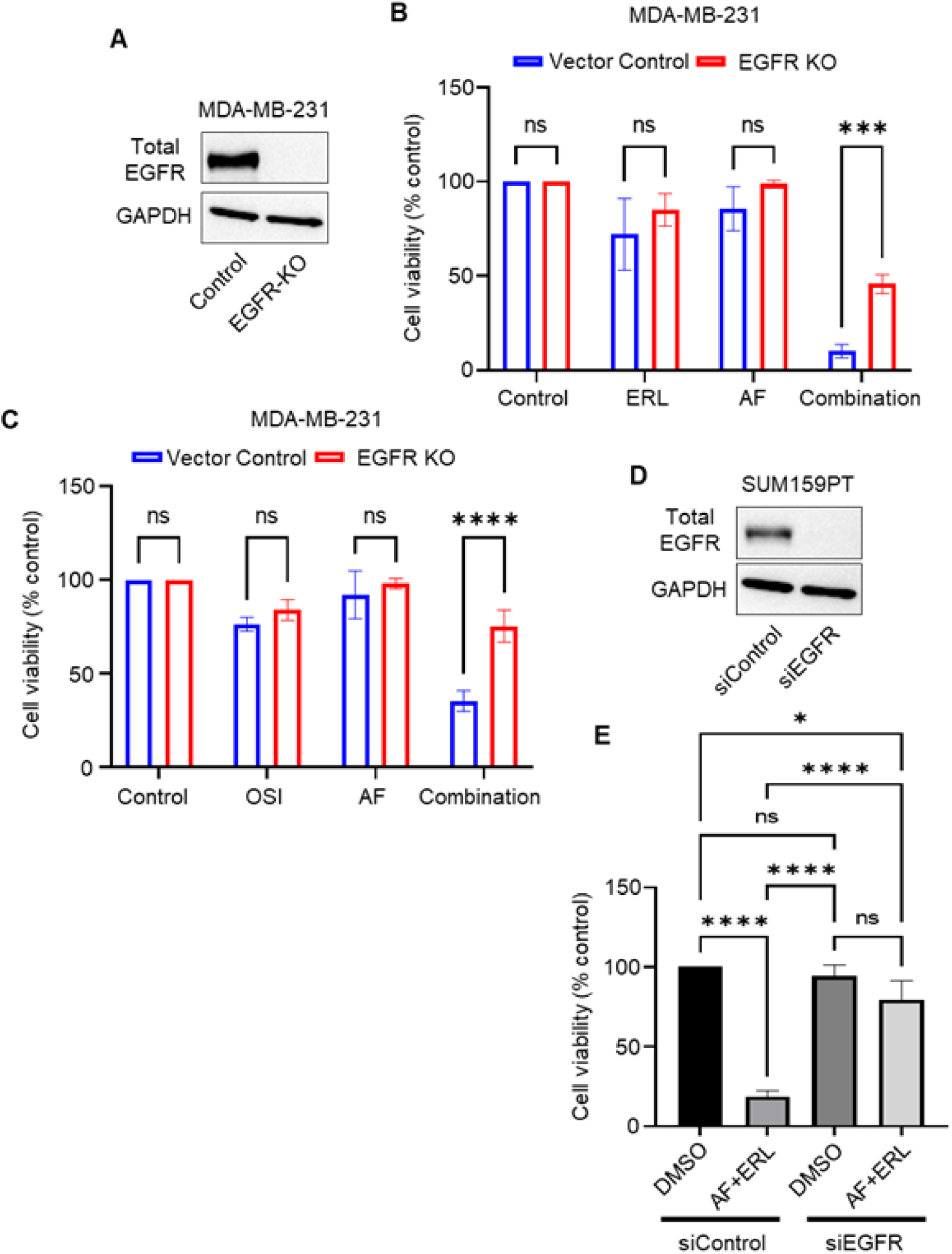
EGFR depletion rescues EGFR-high breast cancer cells from undergoing cell death upon Auranofin-EGFR inhibitor combination treatment. **(A)** Western blot confirming EGFR depletion in CRISPR/Cas9 EGFR KO MDA-MB-231 cells compared to the CRISPR/Cas9 control MDA-MB-231 cells. **(B)** CRISPR/Cas9 control and EGFR KO MDA-MB-231 cells were treated with Auranofin (2.5 µM), Erlotinib (10 µM), either alone or in combination for 72 hours. Cell viability was analyzed by MT Cell viability assays. One-way ANOVA followed by Tukey’s post-test, n = 3 (mean ± SEM). **(C)** CRISPR/Cas9 control and EGFR KO MDA-MB-231 cells were treated with Auranofin (2.5 µM), Osimertinib (5 µM), either alone or in combination for 72 hours. Cell viability was analyzed by MT Cell viability assays. One-way ANOVA followed by Tukey’s post-test, n = 3 (mean ± SEM). **(D)** SUM159PT cells were transfected either with scramble siRNAs (siControl) or EGFR-specific siRNAs for 48 hours. EGFR protein levels were analyzed by Western blot analysis. Representative images of three independent experiments are shown. **(E)** SUM159PT cells transfected with either scramble siRNAs or EGFR-specific siRNAs were treated with Auranofin (2.5 µM), Erlotinib (10 µM), either alone or in combination for 72 hours. Cell viability was analyzed by Crystal Violet cell viability assays. One-way ANOVA followed by Tukey’s post-test, n = 3 (mean ± SEM).

Auranofin is known to inhibit both TXNRD1 and TXNRD3 in cancer cells (22,24). To determine the involvement of TXNRD1 in sensitizing TNBC cells to EGFR inhibitors, we used TRi-1, a specific TXNRD1 inhibitor that selectively suppresses TXNRD1 redox activity (25,26). Interestingly, while TRi-1 significantly inhibited TXNRD1 redox activity in SUM159PT and MDA-MB-231 cells (Fig. S4A), it failed to sensitize these cells to Erlotinib (Fig. S4B). These findings suggest that Auranofin sensitizes TNBC cells to EGFR inhibitors via TXNRD3 inhibition.

### TXNRD3 depletion or inhibition increases EGFR phosphorylation and surface localization in TNBC cells

The presence of activating phosphorylation of EGFR (Y1068) has been linked to better responses to EGFR inhibitors in lung cancer patients (27). Therefore, we examined the expression levels of phosphorylated EGFR protein in TNBC patient tumors using a TNBC tissue microarray. Our results showed that a significant proportion of TNBC tumor cells expressed total EGFR protein (both on the surface and in the cytosol) but completely lacked phosphorylated EGFR protein expression (Fig. S5). This may explain the poor response rates of TNBC patients to EGFR inhibitors despite the high levels of total EGFR protein.

There is significant evidence suggesting that redox-dependent regulation plays a role in the activation of the EGFR signaling pathway in cancers. The intracellular kinase domain of EGFR contains six cysteine residues, one of which (Cys797) is located in the ATP-binding pocket. Oxidation of Cys797 increases EGFR kinase activity and enhances EGFR phosphorylation (Y1068), leading to the activation of the EGFR signaling pathway (28). Since TXNRD3 is involved in regulating intracellular redox homeostasis (22), we first tested whether siRNA-mediated TXNRD3 knockdown increases intracellular ROS levels in TNBC cells. Our results showed that TXNRD3 knockdown, but not TXNRD1 knockdown, significantly increased intracellular ROS levels in both SUM159PT and MDA-MB-231 cells (Fig. 5A). Next, we examined whether TXNRD3 knockdown-induced ROS increases EGFR activating phosphorylation (Y1068) in TNBC cells. Our results demonstrated that TXNRD3 knockdown markedly increased EGFR phosphorylation (Y1068) in both SUM159PT and MDA-MB-231 cells (Fig. 5B). However, TXNRD1 knockdown had only a marginal effect on increasing EGFR phosphorylation (Y1068) in both cell lines (Fig. 5B). Additionally, TXNRD3 knockdown increased ERK1/2 phosphorylation, a downstream target of EGFR, in both SUM159PT and MDA-MB-231 cells (Fig. S6A), suggesting that TXNRD3 knockdown activates the EGFR signaling pathway in TNBC cells. To further confirm the involvement of ROS in the increased EGFR phosphorylation observed upon TXNRD3 knockdown, we treated TXNRD3-knockdown SUM159PT cells with N-acetylcysteine (NAC) for 16 hours. Our results showed that NAC treatment markedly reduced EGFR phosphorylation (Y1068) in TXNRD3-depleted SUM159PT cells (Fig. 5C). These findings suggest that TXNRD3 depletion increases EGFR phosphorylation in a ROS-dependent manner.

**Figure 5:**
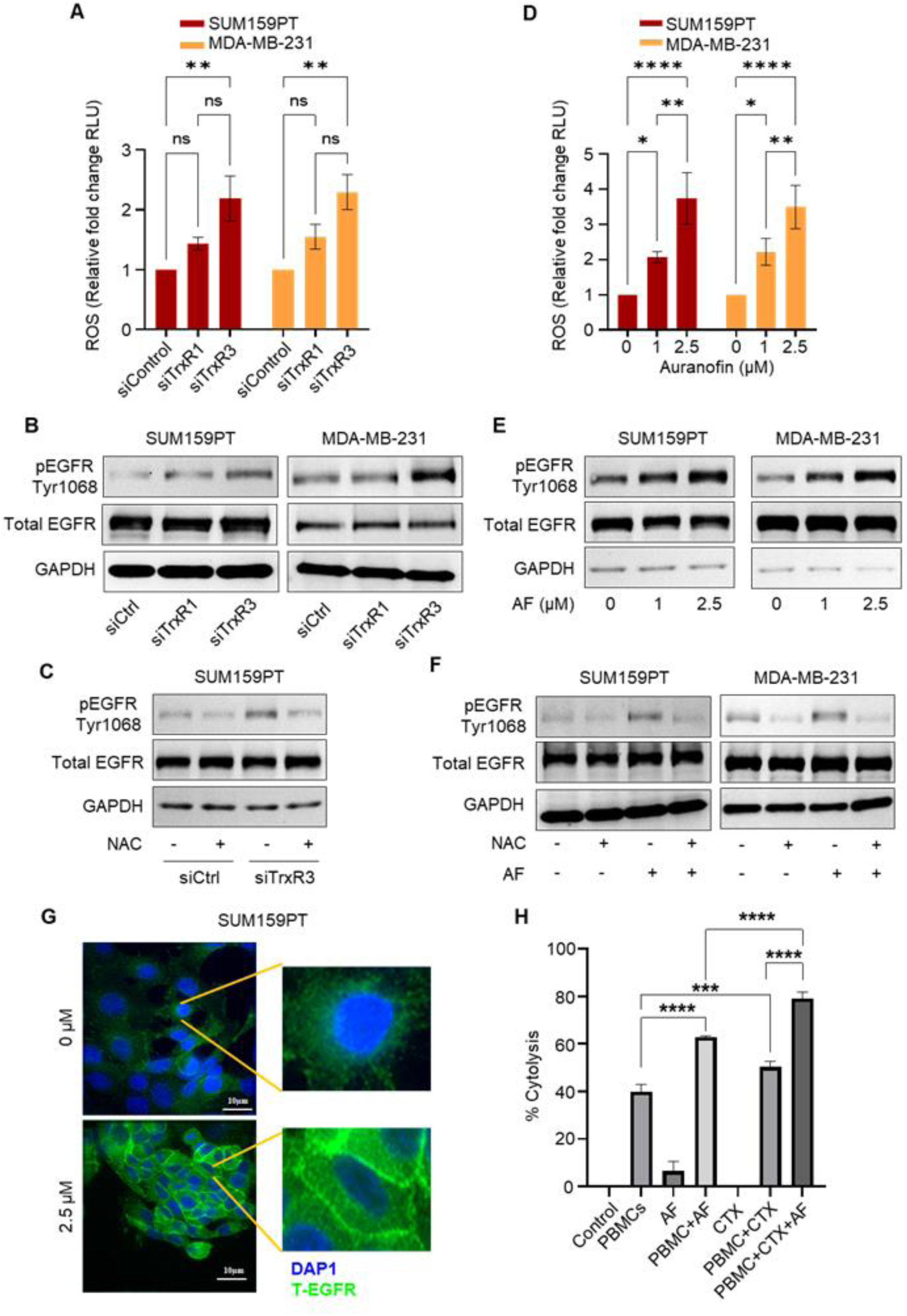
TXNRD3 depletion or inhibition increases EGFR phosphorylation and surface localization in TNBC cells. **(A)** SUM159PT and MDA-MB-231 cells were transfected with either control siRNAs, TrxR1-specific siRNAs, or TrxR3-specific siRNAs for 48 hours, and intracellular ROS levels were analyzed. One-way ANOVA followed by Tukey’s post-test, n = 3 (mean ± SEM). **(B)** SUM159PT and MDA-MB-231 cells were transfected with TrxR1- or TrxR3-specific siRNAs as described above. Protein levels of phospho-EGFR (Y1068) and total EGFR were analyzed by Western blot analysis. Representative images of three independent experiments are shown. GAPDH was used as a loading control. **(C)** SUM159PT and MDA-MB-231 cells were transfected with either TrxR1- or TrxR3-specific siRNAs for 16 hours. Cells were then treated with 10 mM N-acetyl Cysteine (NAC) for 36 hours. Protein levels of phosphor-EGFR (Y1068) and total EGFR were analyzed by Western blot analysis (n=3). **(D)** SUM159PT and MDA-MB-231 cells were treated with Auranofin (0-2.5 µM) for 24 hours, and intracellular ROS levels were analyzed. One-way ANOVA followed by Tukey’s post-test, n = 3 (mean ± SEM). **(E)** SUM159PT and MDA-MB-231 cells were treated with Auranofin (0-2.5 µM) for 24 hours. Protein levels of phospho-EGFR (Y1068) and total EGFR were analyzed by Western blot analysis (n=3). **(F)** SUM159PT and MDA-MB-231 cells were treated with 10 mM NAC for 3 hours followed by the treatment auth Auranofin (0-2.5 µM) for 24 hours. Protein levels of phospho-EGFR (Y1068) and total EGFR were analyzed by Western blot analysis (n=3). **(G)** SUM159PT cells were treated with Auranofin (2.5 µM) for 24 hours. Surface localization of EGFR was analyzed by immunofluorescence analysis. Representative images of three independent experiments are shown. **(H)** SUM159PT cells were treated with Auranofin (1 µM) or cetuximab (50 µg/mL) alone or in the presence of PBMCs for up to 72 hours. ADCC was measured by analyzing the percentage of cytolysis using xCelligence. One-way ANOVA followed by Tukey’s post-test, n = 3 (mean ± SEM).

Next, we examined whether Auranofin also increases EGFR activating phosphorylation (Y1068) in a ROS-dependent manner in TNBC cells. Our results showed that Auranofin significantly increased intracellular ROS levels in SUM159PT and MDA-MB-231 cells in a concentration-dependent manner (Fig. 5D). Additionally, Auranofin treatment markedly increased EGFR phosphorylation (Y1068) in both SUM159PT and MDA-MB-231 cells (Fig. 5E), an effect that was significantly reduced when cells were pre-treated with NAC (Fig. 5F). Auranofin treatment also increased ERK1/2 and AKT phosphorylation in SUM159PT and MDA-MB-231 cells, suggesting that Auranofin activates the EGFR pathway in TNBC cells (Fig. S6B). Taking together, our results indicate that TXNRD3 knockdown or inhibition activates EGFR in a redox-dependent manner.

We then investigated whether Auranofin treatment increases EGFR surface localization in TNBC cells. Several studies have reported that TNBC patient tumors exhibit internalized EGFR expression rather than EGFR localization on the plasma membrane of tumor cells (29–31). The absence of EGFR on the plasma membrane may explain the lack of TNBC patient sensitivity to cetuximab. Moreover, inhibiting Src Family Kinases (SFKs) was shown to reduce nuclear EGFR, increase EGFR accumulation on the plasma membrane, and enhance TNBC cell sensitivity to cetuximab (31). Therefore, we examined whether Auranofin increases EGFR accumulation on the plasma membrane of TNBC cells. SUM159PT cells were treated with 2.5 µM Auranofin for 24 hours, and EGFR surface expression was analyzed by immunofluorescence. Our results showed that Auranofin treatment significantly increased EGFR expression on the plasma membrane (Fig. 5G). Next, we investigated whether Auranofin-induced EGFR surface expression sensitizes SUM159PT cells to cetuximab. Specifically, we tested whether Auranofin enhances cetuximab-induced antibody-dependent cellular cytotoxicity (ADCC) in SUM159PT cells. In cell-based in vitro assays, cetuximab requires the presence of peripheral blood mononuclear cells (PBMCs) to initiate cetuximab-induced cytotoxicity. SUM159PT cells were co-cultured with or without activated PBMCs overnight and subsequently treated with a sub-lethal dose of Auranofin (1 µM), cetuximab (50 µg/mL), or their combination. Our results showed that sub-lethal doses of Auranofin and cetuximab alone did not induce cytolysis in SUM159PT cells in the absence of PBMCs (Fig. 5H). However, we observed approximately 40–55% cytolysis of SUM159PT cells induced by PBMCs alone, PBMCs + Auranofin, and PBMCs + cetuximab (Fig. 5H). Notably, the combination of PBMCs + cetuximab + Auranofin significantly increased SUM159PT cell cytolysis to 80% (Fig. 5H). The percentage of maximal tumor cell death significantly increased (p < 0.01) when comparing cetuximab treatment with and without Auranofin, indicating that Auranofin treatment significantly enhances cetuximab-mediated ADCC (Fig. 5H). Taken together, our findings suggest that Auranofin increases EGFR surface expression and sensitizes TNBC cells to cetuximab.

### Auranofin-Erlotinib combination treatment exerts significant uiuhanti-cancer activity *in vivo*

Next, we evaluated the in vivo anti-cancer efficacy of Auranofin and Erlotinib combination therapy using a human MDA-MB-231 xenograft model. Our results showed that monotherapy with either Auranofin (at half the maximum tolerated dose) or Erlotinib had no significant impact on tumor growth. However, the combination of Auranofin and Erlotinib significantly inhibited MDA-MB-231 tumor growth in vivo (Fig. 6A), indicating enhanced anti-cancer activity in TNBC. The combination therapy was well-tolerated, as no significant weight loss was observed in treated mice compared to controls (Fig. 6B).

**Figure 6:**
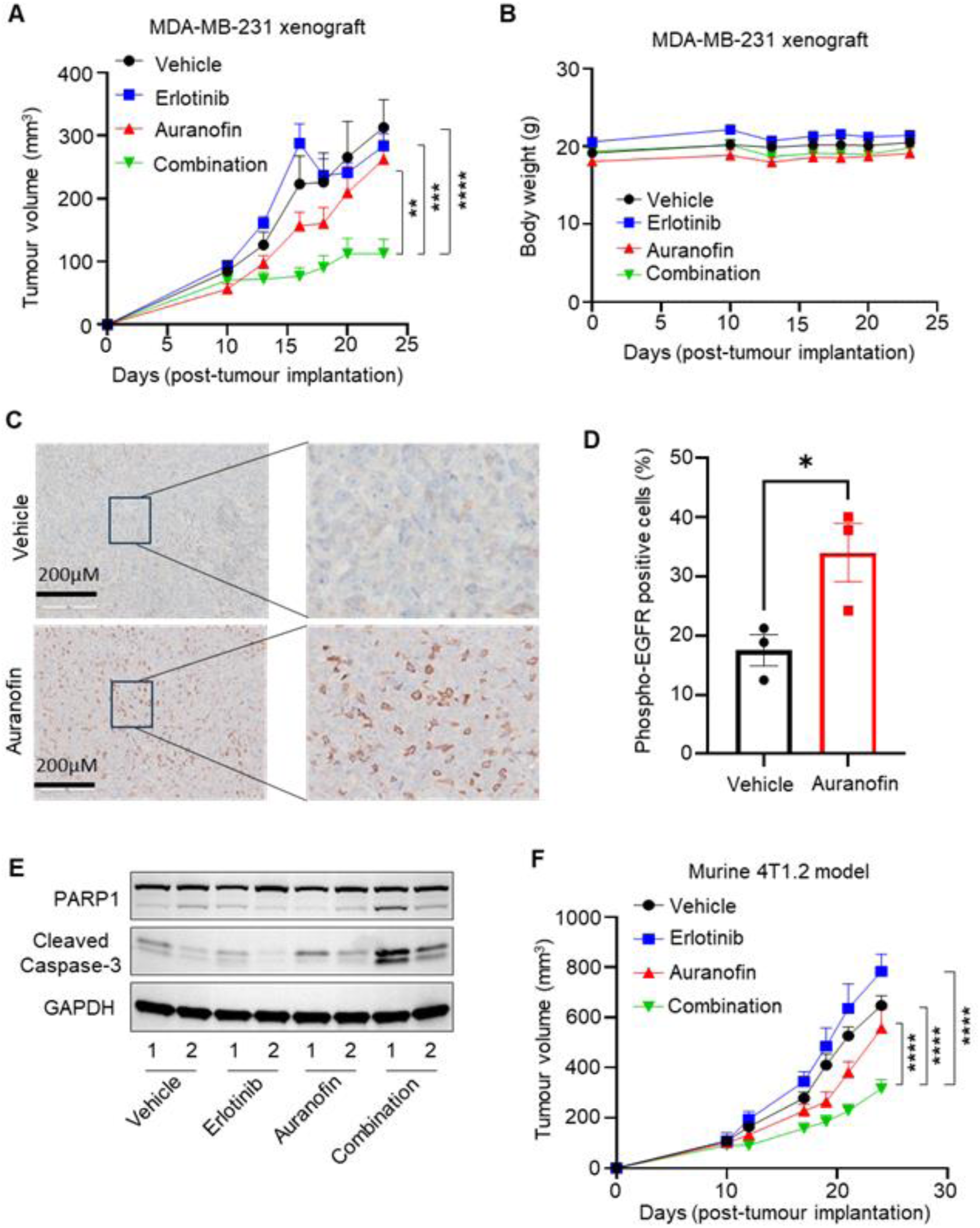
Auranofin-Erlotinib combination treatment exerts significant anti-cancer activity *in vivo*. **(A, B)** Tumor growth (A) and body weight (B) in Balb/c Nude mice orthotopically injected with MDA-MB-231 cells following treatment with vehicle, Erlotinib (50 mg/kg, Monday-Friday, oral gavage), Auranofin (5 mg/kg, Monday-Friday, ip), or combination for two weeks. Treatment started when the tumor reached 50-100 mm^3^. Data are presented as mean ± SEM (n=6 mice/group); Two-way ANOVA followed by Tukey’s post-test were performed on tumor volumes, **p<0.01, ***p<0.001, ****p<0.0001. **(C, D)** Representative IHC images of phospho-EGFR (Y1068) staining of primary MDA-MB-231 tumors treated with vehicle or Auranofin for 2-weeks (C). Quantification of phospho-EGFR positive tumor cells in the primary MDA-MB-231 tumors. Percentage of phospho-EGFR positive tumor cells are presented as mean ± SD (n=3 tumors/group) (D). One-way ANOVA with Sidak’s multiple comparisons test, *p<0.05, **p<0.01. **(E)** Protein levels of cleaved PARP1 and cleaved caspase-3 in MDA-MB-231 tumors treated with Auranofin or Erlotinib alone or combination were analyzed by Western blot analysis. Two representative tumors were used for each treatment group. **(F)** Tumor growth in Balb/c mice orthotopically injected with murine 4T1.2 TNBC cells following treatment with vehicle, Erlotinib (50 mg/kg, Monday-Friday, oral gavage), Auranofin (5 mg/kg, Monday-Friday, ip), or combination for two weeks. Treatment started when the tumor reached 50-100 mm3. Data are presented as mean ± SEM (n=6 mice/group); Two-way ANOVA followed by Tukey’s post-test were performed on tumor volumes, ****p<0.0001.

To investigate the mechanism underlying this effect, we examined whether Auranofin increased EGFR phosphorylation in MDA-MB-231 tumors in vivo. Immunohistochemical analysis of tumors after two weeks of treatment with either vehicle or Auranofin revealed that Auranofin significantly increased levels of phosphorylated EGFR (Y1068) (Fig. 6C). The percentage of phospho-EGFR-positive cells increased from 17.52% in vehicle-treated tumors to 35% in Auranofin-treated tumors, with a mean difference of 16.47% ± 5.596% (Fig. 6D). These findings suggest that Auranofin sensitizes MDA-MB-231 tumors to Erlotinib by activating EGFR.

We further assessed whether Auranofin and Erlotinib combination therapy induced apoptosis in MDA-MB-231 tumors by evaluating the cleavage of caspase-3 and its substrate, PARP1, both classical markers of apoptosis. Compared to monotherapy or vehicle treatment, the combination of Auranofin and Erlotinib markedly increased caspase-3 and PARP1 cleavage (Fig. 6E), confirming enhanced apoptotic activity.

Additionally, we tested the anti-cancer efficacy of Auranofin-Erlotinib combination therapy in a fully immunocompetent murine syngeneic model. The 4T1.2 cell line, a highly aggressive murine TNBC model, mimics human TNBC phenotypes when implanted orthotopically in the mammary fat pad of immunocompetent Balb/c mice. Our results showed that while monotherapy with Auranofin or Erlotinib had no significant effect on tumor growth, the combination treatment significantly reduced 4T1.2 tumor progression (Fig. 6F). Collectively, these findings demonstrate that Auranofin-Erlotinib combination therapy effectively inhibits TNBC tumor growth in vivo, supporting its potential as a therapeutic strategy for TNBC.

## DISCUSSION

TNBCs are commonly characterized by EGFR overexpression (5,6) which has led to several clinical trials evaluating the efficacy of small-molecule EGFR inhibitors and anti-EGFR monoclonal antibodies. However, the results have been disappointing for EGFR inhibitors, both as monotherapy and in combination with standard chemotherapy treatments. Phase II clinical trials of gefitinib and Erlotinib monotherapy showed partial responses of only 0-3% in metastatic and recurrent breast cancer patients (10,32). Erlotinib was also tested in combination with the chemotherapy drug Bendamustine in late-stage TNBC patients, but no clinical efficacy was observed (33). A second-generation EGFR inhibitor afatinib was tested in metastatic TNBC patients and no objective response was observed in these patients (34). In addition to small-molecule inhibitors, the anti-EGFR monoclonal antibody Cetuximab has also been evaluated in multiple clinical trials, either as monotherapy or in combination with chemotherapy, in metastatic breast cancer patients (11) (35) (36). Cetuximab was tested in combination with carboplatin in metastatic recurrent breast cancer patients, showing a 6% response rate with Cetuximab alone and 16% with the combination therapy. However, no statistically significant difference in partial response was observed compared to chemotherapy alone (11). Another clinical trial reported no significant difference in response rates between Cetuximab alone and Cetuximab-cisplatin combination therapy in advanced TNBC patients (35). Interestingly, a Phase II clinical trial for the combination of Cetuximab and ixabepilone showed some clinical efficacy in TNBC patients compared to non-TNBC patients (36). Overall, despite EGFR overexpression, advanced TNBC patients show poor responses to EGFR inhibitors and anti-EGFR monoclonal antibodies. Several mechanisms of resistance, such as network rewiring through the overexpression of MET, HER3, and AXL, have been proposed, but their therapeutic impact requires further evaluation (37). Thus, understanding why TNBCs are intrinsically resistant to EGFR-targeted therapies, despite EGFR overexpression, has become a critical clinical concern.

Genome-wide CRISPR/Cas9 knock-out library screening has enabled researchers to identify the biomarkers of response for a given targeted therapy. Using GeCKO v2A library, which has previously been used for identifying druggable targets in cancer cells (38,39), we found that loss of a redox protein thioredoxin reductase 3 sensitized EGFR-high MDA-MB-231 cells. Functionally, we validated the effect of TXNRD3 loss in EGFR inhibitor sensitization by using TXNRD3-specific siRNAs, where TXNRD3 knocked down significantly sensitized TNBC cells to Erlotinib (first-generation EGFR inhibitor) and Osimertinib (third-generation EGFR inhibitor). Our CRISPR-based knock-out library screening results also echoed with previous study that identified TXNRD3 as a key molecule in conferring sorafenib, a multi-kinase inhibitor, resistance in leukemia cells (22).

The expression and redox activity of thioredoxin (Trx) and thioredoxin reductase (TXNRD) are upregulated in various cancers, including TNBCs (24,40,41) and are associated with drug resistance (42). Amongst the three isoforms of TXNRD, TXNRD1 is located in the cytosol, TXNRD2 in mitochondria (43), and TXNRD3 in both cytosol (66.6 kDa) and mitochondria (70.7 kDa) (22). We previously demonstrated that TXNRD1 is upregulated in TNBC cells, and its inhibition using the FDA-approved inhibitor Auranofin significantly reduced TNBC tumor growth (24). A recent study also showed that TXNRD3 expression is notably higher in TNBC tumor tissues compared to non-TNBC tissues and normal breast epithelial cells (23). Consistent with this, we found that TXNRD3 expression, but not TXNRD1, correlated with resistance to EGFR inhibitors in TNBC cells. Depletion of TXNRD3, but not TXNRD1, sensitized TNBC cells to EGFR inhibitors. Mechanistically, we observed a significantly higher accumulation of reactive oxygen species (ROS) upon depletion of TXNRD3 (the mitochondrial isoform) compared to TXNRD1. Additionally, a TXNRD1-specific inhibitor, TRi-1, (25,26), significantly reduced TXNRD1 redox activity in TNBC cells but did not sensitize these cells to Erlotinib or Osimertinib.

We also used Auranofin, an FDA-approved thioredoxin reductase inhibitor that targets both TXNRD isoforms (22) (19). Our results showed that a sub-lethal dose of Auranofin combined with either Erlotinib or Osimertinib exerted synergistic anti-cancer activity in vitro in EGFR-high TNBC cells. Moreover, Auranofin treatment significantly enhanced the anti-cancer activity of Erlotinib in MDA-MB-231 cell line-derived xenografts as well as in a murine syngeneic 4T1.2 model of TNBC in vivo. In this study, we demonstrated that Auranofin significantly reduced TXNRD3 levels in TNBC cells, indicating that TXNRD3 inhibition sensitizes TNBC cells to EGFR inhibitors.

A key question is why TNBC cells, despite having high EGFR levels, are resistant to EGFR-targeted therapies, and how TXNRD3 inhibition alters their sensitivity to EGFR inhibitors. Traditionally, EGFR is localized on the plasma membrane, functioning as a receptor tyrosine kinase to promote cancer cell proliferation (44). However, recent studies have shown that EGFR can also be localized in intracellular organelles, including the nucleus and cytosol (31,45). Increased intracellular EGFR is associated with resistance to cetuximab in non-small cell lung cancer (46,47) and TNBCs (31). Therefore, the lack of EGFR on the plasma membrane in TNBC cells may explain the poor response to Cetuximab in these patients. Furthermore, EGFR activation can be modulated via cysteine oxidation or sulfenylation. Cysteine 797 in the catalytic domain of EGFR undergoes sulfenylation in response to elevated intracellular ROS, leading to increased EGFR activation or kinase activity (28). Consistent with these findings, we observed that both TXNRD3 knockdown and Auranofin treatment significantly increased intracellular ROS levels and EGFR phosphorylation (Y1068) in SUM159PT and MDA-MB-231 cells. Pre-treatment of these cells with N-acetylcysteine, a ROS scavenger, markedly reduced EGFR phosphorylation (Y1068) following TXNRD3 knockdown and Auranofin treatment. These results suggest that TXNRD3 depletion or inhibition activates EGFR in a redox-dependent manner.

Phosphorylated EGFR (Y1068) is an active form of protein, and increased levels of phosphorylated EGFR have been shown to correlate with a better response to Erlotinib in non-small cell lung cancer patients (27). In conclusion, our study identified elevated TXNRD3 expression as a genetic marker of intrinsic resistance to EGFR inhibitors in TNBCs, using unbiased whole-genome CRISPR-library screening. This study presents a novel therapeutic strategy, using Auranofin to modulate EGFR activity in a redox-dependent manner, and sensitizing EGFR-high TNBCs to EGFR inhibitors. Our findings provide a strong rationale for translating Auranofin and EGFR inhibitor combination therapy for EGFR-high TNBC patients. However, for clinical translation, robust markers of response to this combination therapy must be identified to stratify patients who are most likely to benefit from this targeted approach.

## Supporting information

Supplemental CRISPR screening data

## ACKNOWLEDGEMENT

We thank the personnel of the QIMRB and TRI Animal facilities for their assistance in housing and husbandries of animals during the experiments. We also thank Mater Pathology for providing breast cancer patient tissue microarray slides. We also thank QIMRB Histology facility for staining the immunohistochemistry slides.

## AUTHOR CONTRIBUTION

PR contributed to study concept and design, acquisition of data, analysis, and interpretation of data. PR, GG, MH, MK, and KKK contributed to experimental design, data analysis, interpretation of data, and in writing of the manuscript. PR carried out all *in vitro* work and animal experiments presented in this manuscript. GG performed bioinformatic analysis of the CRISPR library screening data. MH provided intellectual input in designing CRISPR screening and data analysis. SS performed immunofluorescence experiment and data analysis. GA performed ADCC experiment and analyzed the data. All authors contributed to the review and revision of this manuscript.

## DATA AVAILABILITY

The datasets used and/or analyzed during the current study are available from the corresponding author on reasonable request. The materials used in this study are available from the corresponding author upon request. The whole genome sequencing data associated with the CRISPR library screening experiment has been submitted to GEO under the metadata file name “240910-GEO-ERL-CRISPR-RNAseq.xlsx”.

