## Supplementary materials, methods, and figures for "Thioredoxin reductase 3 inhibition sensitizes triple-negative breast cancer cells to EGFR inhibitors"

**Table S1: List of antibodies.**

| Marker | Cat # | Titration | Vendor |
|---|---|---|---|
| TrxR1 | MAB7428 | 1:1000 | R&D Systems |
| TrxR3 | abx239135 | 1:1000 | Abbexa |
| Phospho-EGFR (Y1068) | 2234S | 1:1000 | Cell Signaling |
| Total EGFR | 4267T | 1:1000 | Cell Signaling |
| Vinculin | 13901S | 1:1000 | Cell Signaling |
| Phospho-ERK1/2 | | 1:1000 | Cell Signaling |
| Total ERK1/2 | | 1:1000 | Cell Signaling |
| Phospho-AKT1 | | 1:1000 | Cell Signaling |
| Total AKT1 | | 1:1000 | Cell Signaling |
| GAPDH | RDS2275PC100 | 1:1000 | R&D Systems |

**Table S2: List of siRNAs.**

| Gene | siRNA Sequence (Sense) | siRNA Sequence (Anti-sense) |
|---|---|---|
| EGFR | GGAUAUUCUGAAAACCGUAAAGGAA | GACCUAUAAGACUUUUGGCAUUUCCUU |
| TXNRD1 | ACAAGUACAUCUGCGAUCAACUCTA | UAGAGUUGAUCGCAGAUGUACUUGUUU |
| TXNRD3 | UGAUAACCUUGAAGCUAUUCUCCCC | CCACUAUUGGAACUUCGAUAAGAGGGG |

**SUPPLEMENTARY FIGURES:**

**Figure S1:**

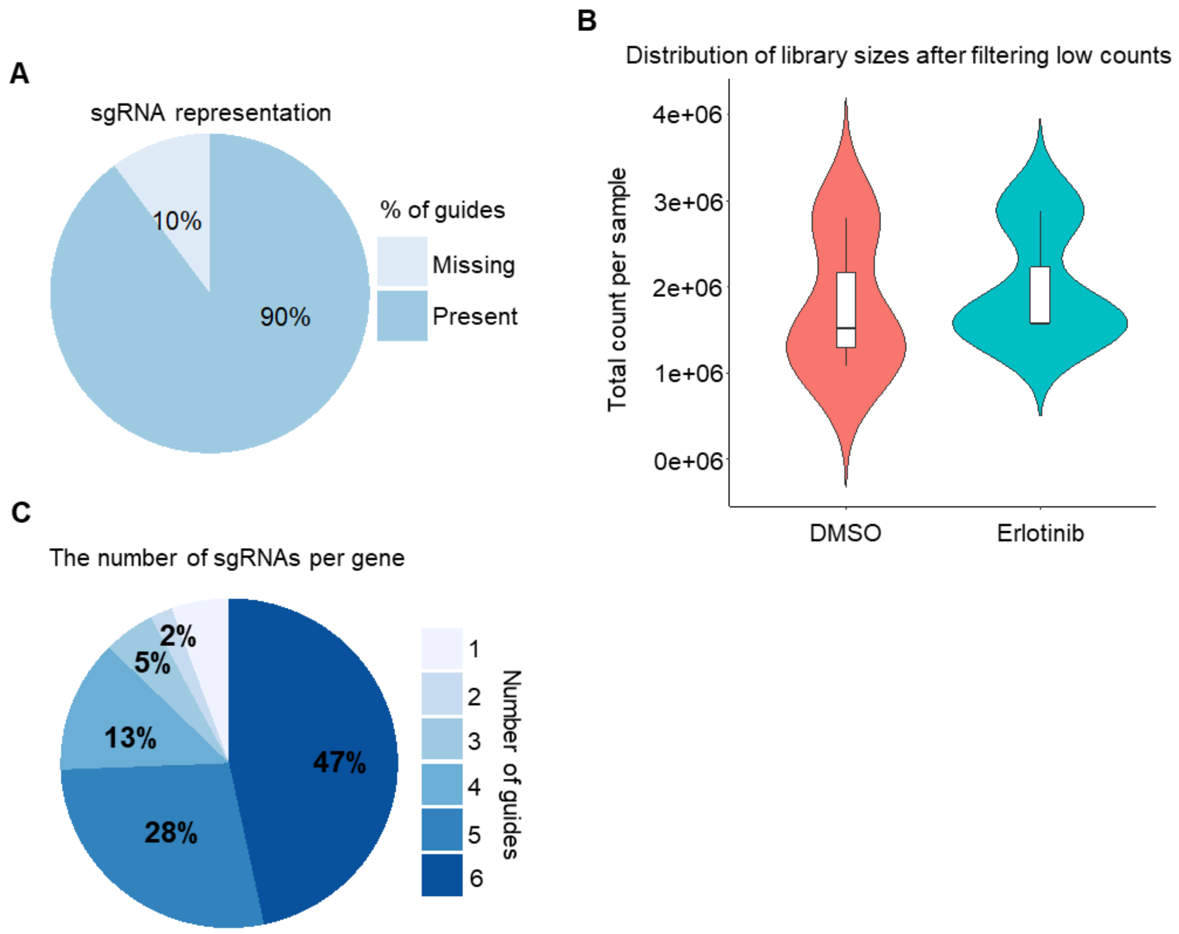


**Figure S1: Genome wide CRISPR screening using GeCKO v2 library in MDA-MB-231 cells.**

**(A)** The pie chart showing the percentage of sgRNAs missing and present in MDA-MB-231 cells after transduction of the GeCKO v2 library and puromycin selection.

**(B)** The pie chart representing the number of guides per gene. The chart also displays the percentage of genes targeted by a specific number of sgRNAs in our genome wide CRISPR screening using GeCKO v2 library.

**(C)** The violin plot of the distribution of GeCKO v2 library. The figure illustrates the distribution of the sgRNA abundances in each library depicting total counts per condition in cells treated with DMSO and Erlotinib. These counts reflect the filtering of non-targeting sgRNAs and sgRNAs with low expression.

**Figure S2:**

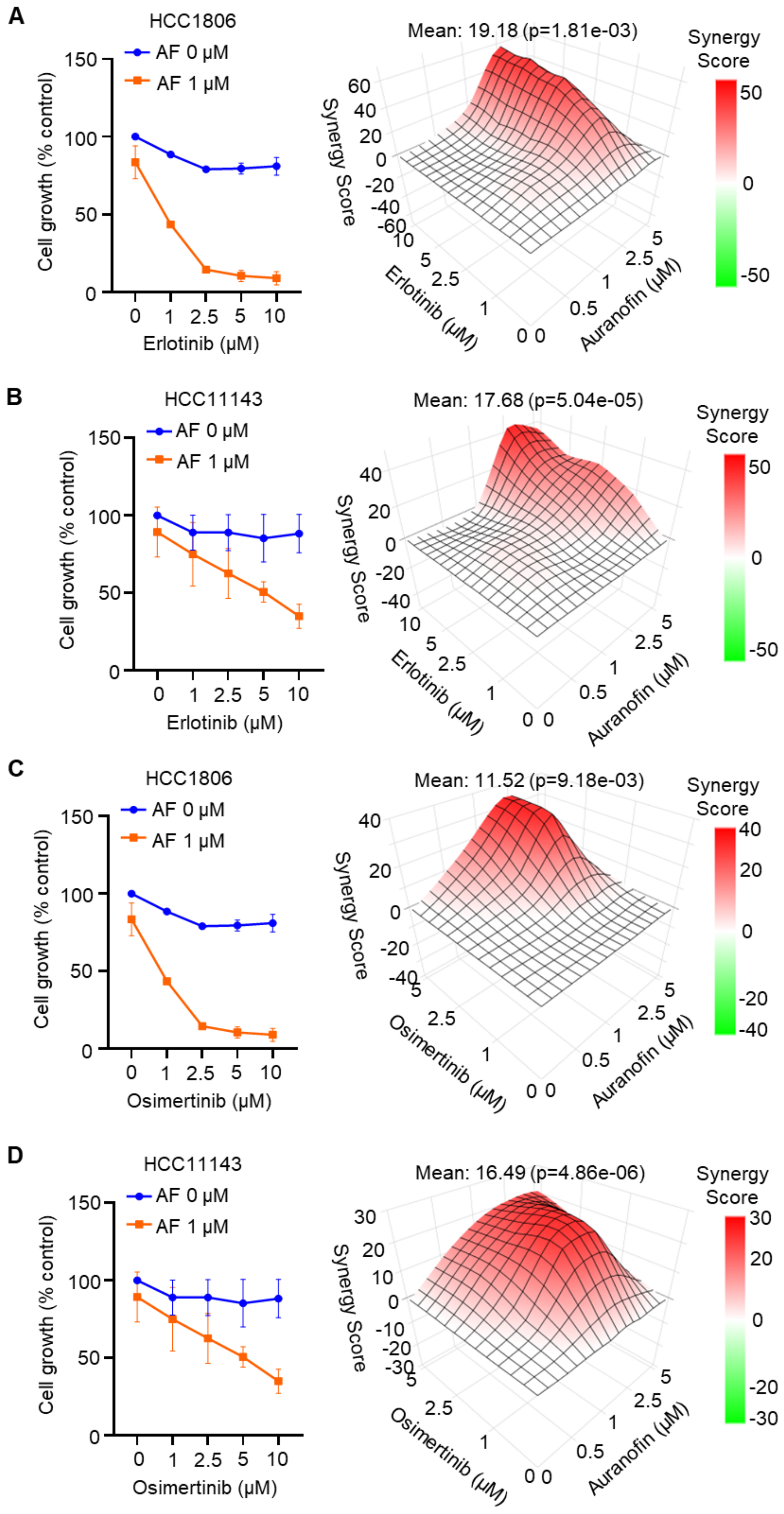
A
HCC1806
AF 0 μM
AF 1 μM
Cell growth (% control)
Erlotinib (μM)
Mean: 19.18 (p=1.81e-03)
Synergy Score
Erlotinib (μM)
Auranofin (μM)
B
HCC11143
AF 0 μM
AF 1 μM
Cell growth (% control)
Erlotinib (μM)
Mean: 17.68 (p=5.04e-05)
Synergy Score
Erlotinib (μM)
Auranofin (μM)
C
HCC1806
AF 0 μM
AF 1 μM
Cell growth (% control)
Osimertinib (μM)
Mean: 11.52 (p=9.18e-03)
Synergy Score
Osimertinib (μM)
Auranofin (μM)
D
HCC11143
AF 0 μM
AF 1 μM
Cell growth (% control)
Osimertinib (μM)
Mean: 16.49 (p=4.86e-06)
Synergy Score
Osimertinib (μM)
Auranofin (μM)

**<u>Figure S2:</u> Pharmacological inhibition of TXNRD3 using auranofin sensitized TNBC cells to EGFR inhibitors *in vitro*.**

**(A, B)** HCC1806 (A) and HCC1143 (B) cells were treated with auranofin (AF) (0-5 μM) and Erlotinib (0-10 μM), both alone and in combination, for 72 hours, and cell viability was analyzed by MT cell viability assays. For cell growth curves (left panel), results with only 1 μM AF are shown. Synergy score was calculated using Synergy Finder. One-way ANOVA followed by Tukey's post-test, n = 3 (mean ± SEM).

**(C, D)** HCC1806 (C) and HCC1143 (D) cells were treated with auranofin (0-5 μM) and Osimertinib (0-10 μM), both alone and in combination, for 72 hours, and cell viability was analyzed by MT cell viability assays. For cell growth curves (left panel), results with only 1 μM AF are shown. Synergy score was calculated using Synergy Finder. One-way ANOVA followed by Tukey's post-test, n = 3 (mean ± SEM).

**Figure S3:**

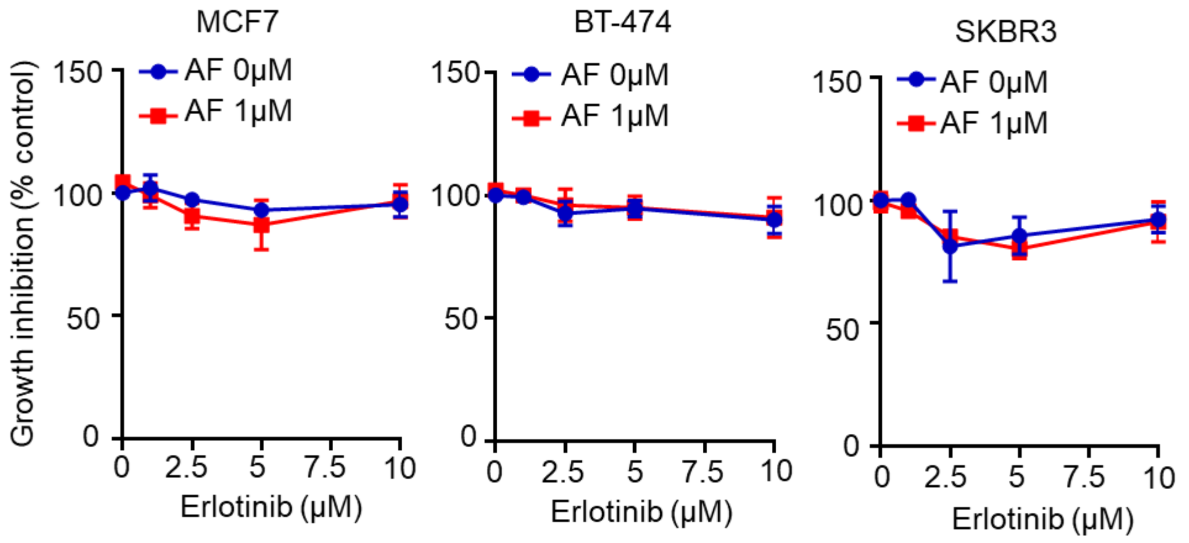


**Figure S3: Effect of auranofin on sensitizing EGFR-low breast cancer cells to Erlotinib *in vitro*.**

EGFR-low non-TNBC cell lines including MCF7, BT-474, and SKBR3 were treated Erlotinib (0-10 μM) alone or in combination with auranofin (AF) (1 μM) for 72 hours. Cell viability was analyzed by MT cell viability assay. One-way ANOVA followed by Tukey's post-test, n = 3 (mean ± SEM).

**Figure S4:**

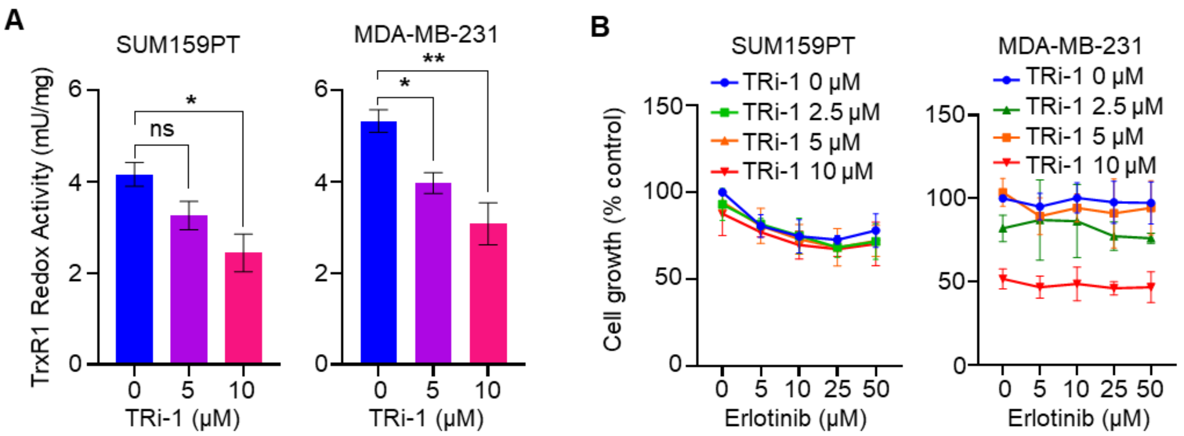


**Figure S4: Effect of thioredoxin reductase 1 specific inhibitor, TRi-1, on sensitizing TNBC cells to Erlotinib.**

**(A)** SUM159PT and MDA-MB-231 cells were treated with TRi-1 (0-10 μM) for 24 hours. The redox activity of thioredoxin reductase 1 (TrxR1) was analyzed by the DTNB reduction assays. One-way ANOVA followed by Tukey's post-test, n = 3 (mean ± SEM).

**(B)** SUM159PT and MDA-MB-231 cells were treated with TRi-1 (0-10 μM) alone or in combination with Erlotinib (0-50 μM) for 72 hours. Cell viability was analyzed by MT cell viability assay. One-way ANOVA followed by Tukey's post-test, n = 3 (mean ± SEM).

**Figure S5:**

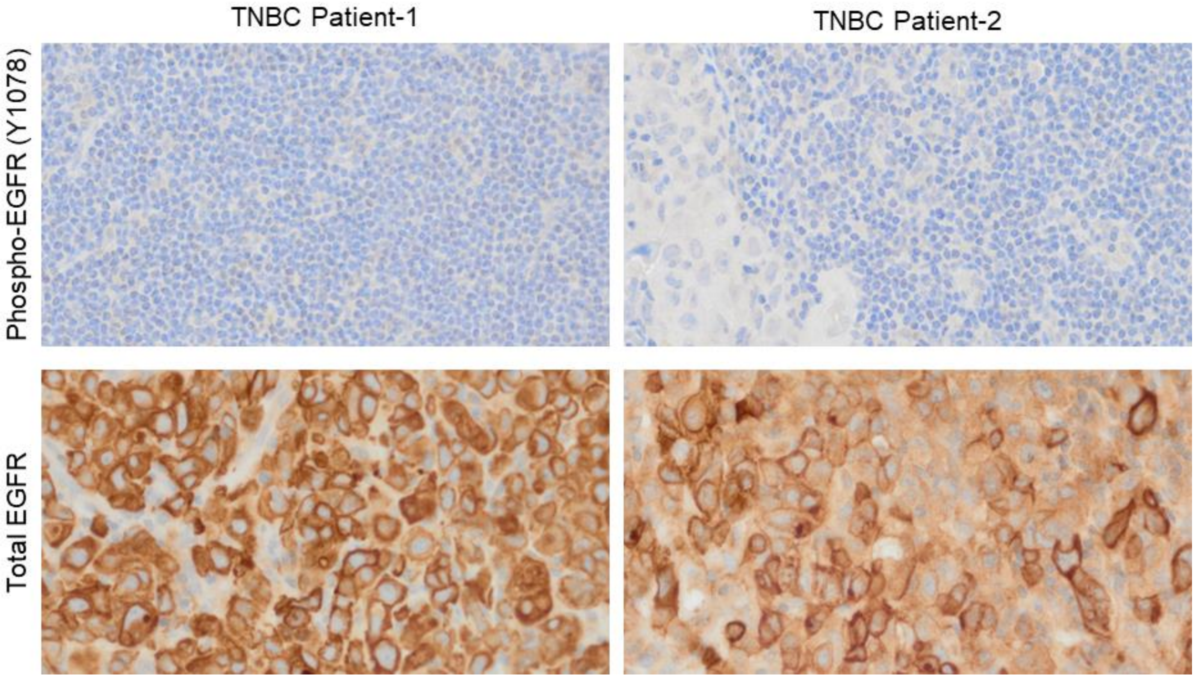


**Figure S5: Expression of Phospho-EGFR and total EGFR protein on TNBC patient tumour cells.**

Tissue microarray analysis of primary and treatment naïve TNBC patient tumours for the expression of phosphorylated EGFR (Y1078) protein and total EGFR protein. Representative images from two TNBC patient tumours are shown.

**Figure S6:**

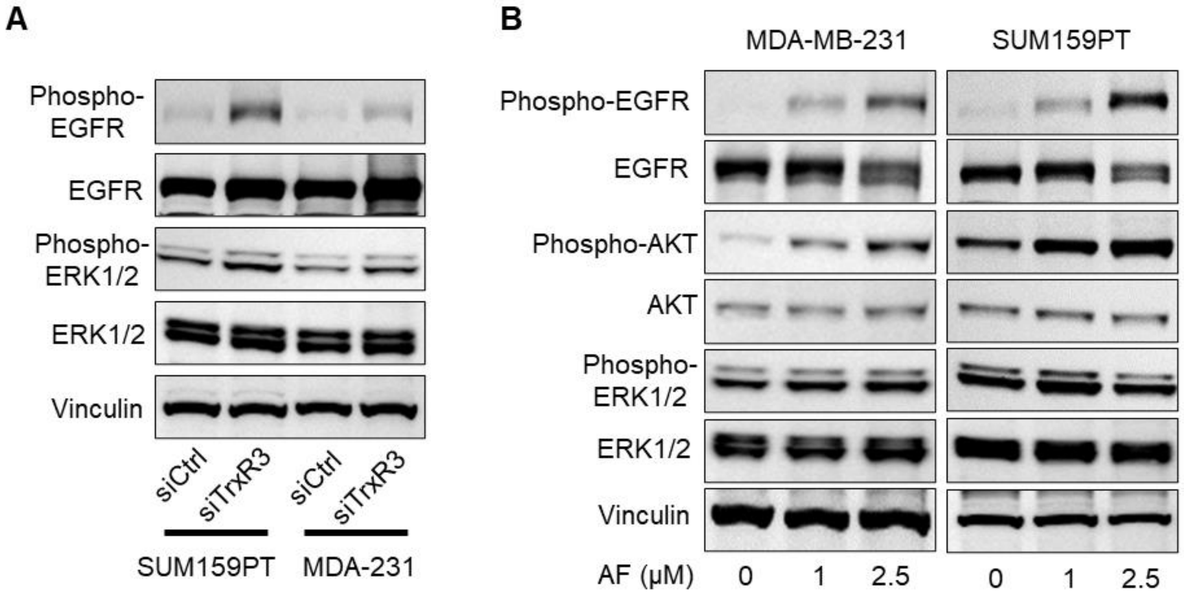


**Figure S6: TXNRD3 knockdown and inhibition activates EGFR signalling pathway in TNBC cells.**

**(A)** SUM159PT and MDA-MB-231 cells were transfected with either control siRNAs, TrxR1-specific siRNAs, or TrxR3-specific siRNAs for 48 hours. Protein levels of phospho-ERK1/2 and total ERK1/2 were analyzed by Western blot analysis. Representative images of three independent experiments are shown. Vinculin was used as a loading control.

**(B)** SUM159PT and MDA-MB-231 cells were treated with auranofin (0-2.5 µM) for 24 hours. Protein levels of phospho-ERK1/2, total ERK1/2, phospho-AKT, and total AKT were analyzed by Western blot analysis. Representative images of three independent experiments are shown. Vinculin was used as a loading control.
